# Rapid isolation and profiling of a diverse panel of human monoclonal antibodies targeting the SARS-CoV-2 spike protein

**DOI:** 10.1101/2020.05.12.091462

**Authors:** Seth J. Zost, Pavlo Gilchuk, Rita E. Chen, James Brett Case, Joseph X. Reidy, Andrew Trivette, Rachel S. Nargi, Rachel E. Sutton, Naveenchandra Suryadevara, Elaine C. Chen, Elad Binshtein, Swathi Shrihari, Mario Ostrowski, Helen Y. Chu, Jonathan E. Didier, Keith W. MacRenaris, Taylor Jones, Samuel Day, Luke Myers, F. Eun-Hyung Lee, Doan C. Nguyen, Ignacio Sanz, David R. Martinez, Ralph S. Baric, Larissa B. Thackray, Michael S. Diamond, Robert H. Carnahan, James E. Crowe

## Abstract

Antibodies are a principal determinant of immunity for most RNA viruses and have promise to reduce infection or disease during major epidemics. The novel coronavirus SARS-CoV-2 has caused a global pandemic with millions of infections and hundreds of thousands of deaths to date^1,2^. In response, we used a rapid antibody discovery platform to isolate hundreds of human monoclonal antibodies (mAbs) against the SARS-CoV-2 spike (S) protein. We stratify these mAbs into five major classes based on their reactivity to subdomains of S protein as well as their cross-reactivity to SARS-CoV. Many of these mAbs inhibit infection of authentic SARS-CoV-2 virus, with most neutralizing mAbs recognizing the receptor-binding domain (RBD) of S. This work defines sites of vulnerability on SARS-CoV-2 S and demonstrates the speed and robustness of new antibody discovery methodologies.

---

Human mAbs to the viral surface spike (S) glycoprotein mediate immunity to other betacoronaviruses including SARS-CoV^3-7^ and Middle East respiratory syndrome (MERS)^8-17^. Because of this, we and others have hypothesized that human mAbs may have promise for use in prophylaxis, post-exposure prophylaxis, or treatment of SARS-CoV-2 infection^18^. MAbs can neutralize betacoronaviruses by several mechanisms including blocking of attachment of the S protein RBD to a receptor on host cells (which for SARS-CoV and SARS-CoV-2^1^ is angiotensin-converting enzyme 2 [ACE2])^12^. We hypothesized that the SARS-CoV-2 S protein would induce diverse human neutralizing antibodies following natural infection. While antibody discovery usually takes months to years, there is an urgent need to both characterize the human immune response to SARS-CoV-2 infection and to develop potential medical countermeasures. Using Zika virus as a simulated pandemic pathogen and leveraging recent technological advances in synthetic genomics and single-cell sequencing, we recently isolated hundreds of human mAbs from a single B cell suspension and tested them *in vitro* for neutralization and for protection in small animals and nonhuman primates, all within 78 days^19^. Using similar methodologies and further efficiency improvements, we sought to obtain human mAbs rapidly for SARS-CoV-2 from the B cells of some of the first human subjects identified with infection in North America. We used an approach similar to that in our previous technical demonstration with Zika, however, for the SARS-CoV-2 discovery effort we report here we used several different workflows in parallel (**Figure 1, Table S2**), which we completed in an expedited time frame (**Figure 1**).

**Figure 1.**
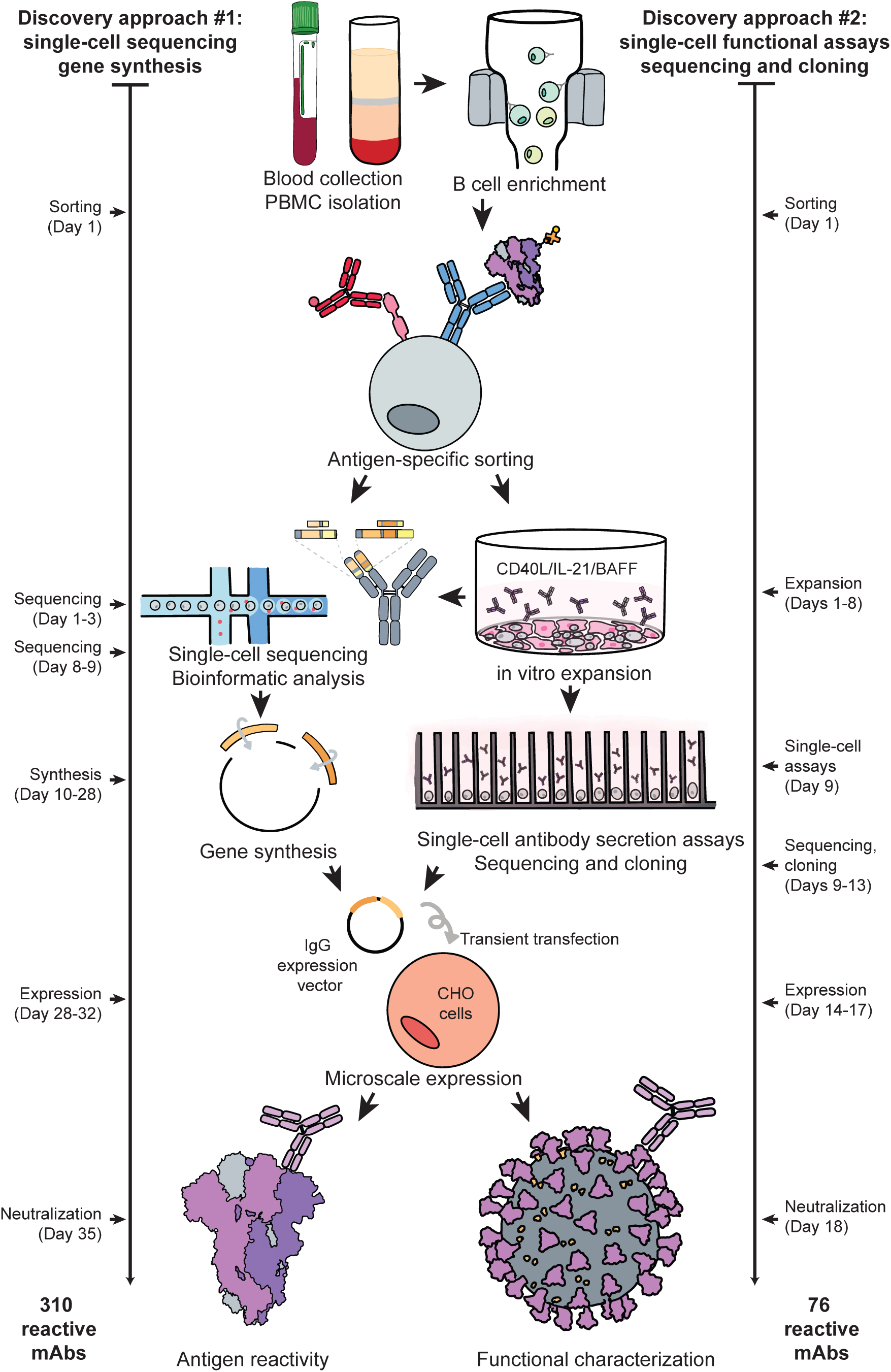
Workflows and timelines. **a. Overview of rapid monoclonal antibody discovery workflows**. The overall scheme is shown, representing the seven specific workflows conducted in parallel (specified in **Table S2**). Blood was collected and white blood cells separated, B cells were enriched from PBMCs by negative selection using paramagnetic beads, antigen-specific cells were obtained by flow cytometric sorting, then processed for direct B cell selection and sequencing or *in vitro* expansion/activation. Cultured B cells were loaded on a Beacon instrument (Berkeley Lights) for functional screening (**Figure S2, Movie S1**) or in a Chromium device (10X Genomics) followed by RT-PCR, sequence analysis, cDNA gene synthesis and cloning into an expression vector, and microscale IgG expression in CHO cells by transient transfection. Recombinant IgG was tested by ELISA for binding to determine antigen reactivity and by a cell impedance-based neutralization test (xCelligence; ACEA) (**Figure S3**) with live virus in a BSL3 laboratory for functional characterization.

We first developed or obtained antigens and recombinant proteins necessary for identifying and isolating antigen-reactive B cells. We synthesized a cDNA encoding a stabilized trimeric prefusion ectodomain of S protein (S2P_ecto_)^20^, expressed the protein in 293F cells, and verified the presence of the prefusion conformation by electron microscopy (**Figure S1)**. We also synthesized and expressed the S protein receptor binding domain (S_RBD_) and obtained recombinant S protein N terminal domain (S_NTD_) that had been prepared by other academic or commercial sources. Using these tools, we designed a mAb discovery approach focused on identifying naturally occurring human mAbs specific for S.

We obtained blood samples from four subjects infected in China who were among the earliest identified SARS-CoV-2-infected patients in North America (**Table S1**). These subjects had a history of recent laboratory-confirmed SARS-CoV-2 infection acquired in Wuhan or Beijing, China. The samples were obtained 35 days (subject 1; the case identified in the U.S.^21^), 36 days (subject 2), or 50 days (subjects 3 and 4) after the onset of symptoms. We tested plasma or serum specimens from the four subjects infected with SARS-CoV-2, or from a healthy donor (subject 5) as control. Serum/plasma antibody ELISA binding assays using S2P_ecto_, S_RBD_, or S_NTD_ protein from SARS-CoV-2 or S2P_ecto_ protein from SARS-CoV revealed that the previously infected subjects had circulating antibodies that recognized each of the proteins tested, with highest reactivity against SARS-CoV-2 S2P_ecto_ and S_RBD_ proteins (**Figure 2a**). Each of the immune subjects also had circulating antibodies that bound to SARS-CoV S2P_ecto_. The healthy donor serum antibodies did not react with any of the antigens. B cells were enriched from PBMCs by negative selection using antibody-coated magnetic beads and stained with phenotyping antibodies specific for CD19, IgD, and IgM. Analytical flow cytometry was performed to assess the frequency of antigen-specific memory B cells for each donor.

**Figure 2.**
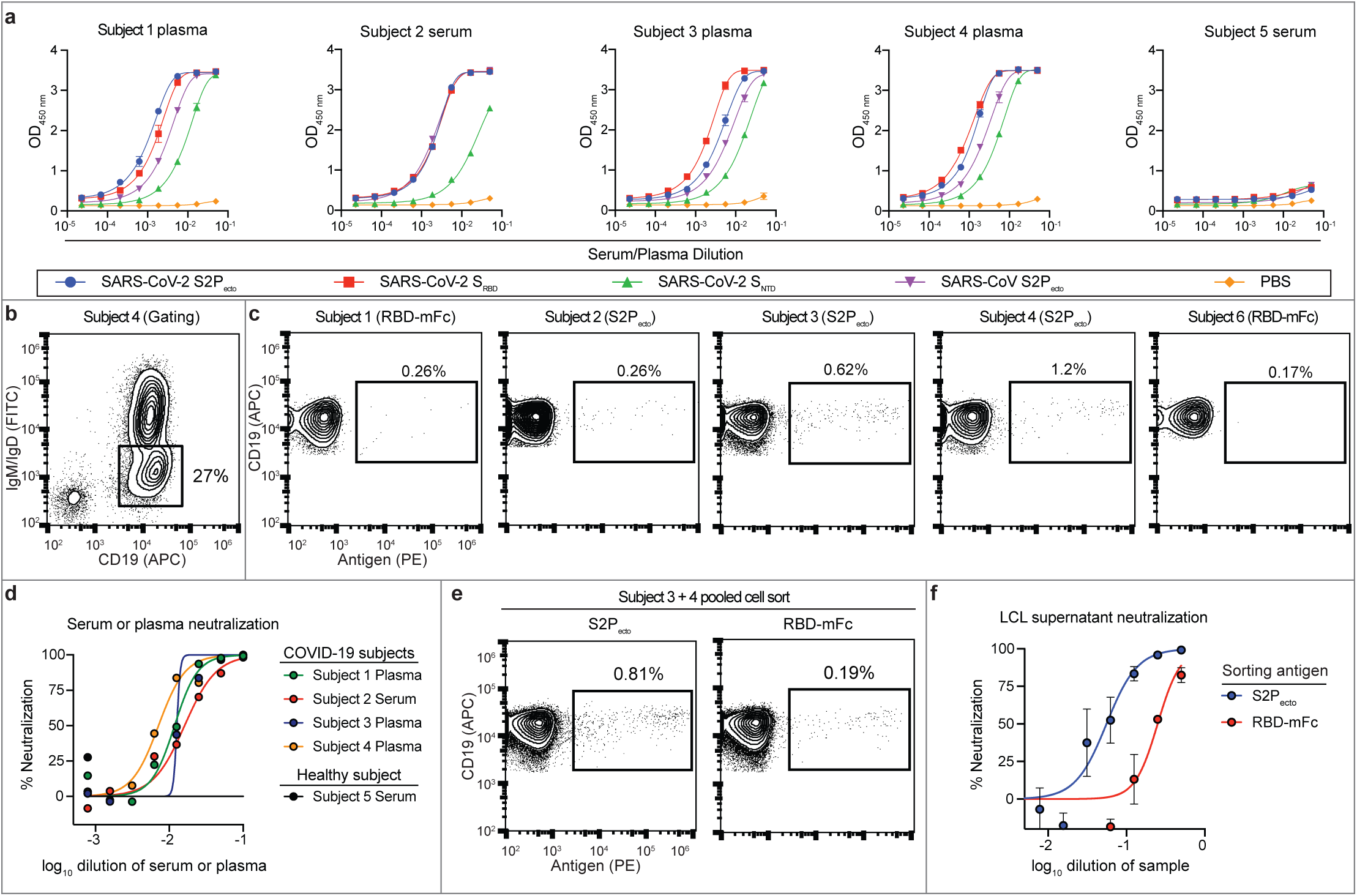
Characterization of SARS-CoV-2 immune donor samples. **a. Serum or plasma antibody reactivity** for the four SARS-CoV-2 immune subjects or one non-immune control, in ELISA using SARS-CoV-2 S2P_ecto_, S_RBD_, S_NTD_, SARS-CoV S2P_ecto_ or PBS. **b. Gating for memory B cells** in total B cells enriched by negative selection using magnetic beads for subject 4; Cells were stained with anti-CD19 antibody conjugated to allophycocyanin (APC) and anti-IgM and anti-IgD antibodies conjugated to fluorescein isothiocyanate (FITC). **c. Analytical flow cytometric analysis of B cells** for subjects 1 to 4, compared to a healthy subject (subject 6). Plots show CD19^+^IgD^-^IgM^-^ population using gating strategy as in **b**. Cells labeled with biotinylated S2P_ecto_ or RBD-mFc antigens were detected using phycoerythrin (PE)-conjugated streptavidin. **d. Plasma or serum neutralizing activity** against the WA1/2020 strain SARS-CoV-2 for subjects 1 to 4 or a healthy donor (subject 6). % neutralization is reported. **e. FACS isolation of** S2P_ecto_ or RBD-mFc-reactive B cells from pooled B cells of subject 3 and 4. Plots show CD19^+^IgD^-^IgM^-^ population using gating strategy as in **b**, and antigen-reactive B cells were identified as in **c**. **f. Lymphoblastoid cell line (LCL) supernatant neutralization**. Neutralization of the WA1/2020 strain SARS-CoV-2 by supernatant collected from cell cultures of S2P_ecto_-or RBD-mFc-sorted memory B cells that had been stimulated in bulk *in vitro* on feeder layers expressing CD40L and secreting IL-21 and BAFF. The supernatants were tested in a ten-point dilution series in the FRNT, and % neutralization is reported. Values shown are the mean ± SD of technical duplicates.

We identified class-switched memory B cells by gating for an IgD^-^/IgM^-^/CD19^+^ population (**Figure 2b)**. From this memory B cell population, we identified antigen-reactive cells using biotinylated recombinant S2P_ecto_ protein or biotinylated RBD fused to mouse Fc (RBD-mFc). Subjects 1 and 2 had very low frequencies of antigen-specific memory B cells that were not greater than two-fold above the background staining frequency in a non-immune sample (subject 6) (**Figure 2c)**. In contrast to subjects 1 and 2, subjects 3 and 4 were 2 weeks later in convalescence and exhibited 0.62 or 1.22 % of class-switched B cells that reacted with antigen (**Figure 2c**). Subjects 3 and 4 also exhibited high titers in a serum antibody focus reduction neutralization test (FRNT) with an authentic SARS-CoV-2 strain (WA/1/2020) (**Figure 2d**). Therefore, we focused subsequent efforts on sorting B cells from the specimens of subjects 3 and 4, which were pooled for efficiency. The pooled memory B cell suspension had frequencies for S2P_ecto_ or RBD-mFc that were 0.81 or 0.19% of the IgD-/IgM-/CD19+ population, respectively (**Figure 2e**). The bulk sorted S2P_ecto_-or RBD-mFc-specific B cells were stimulated on a feeder layer with CD40L, IL-21 and BAFF^22^, and the secreted antibodies in the resulting cell culture supernatants exhibited neutralizing activity against the WA1/2020 strain (**Figure 2f**). After 7 days in culture, these activated B cells were removed from the feeder layers. Roughly half of these B cells were single-cell sequenced and antibody genes were synthesized as previously described^19^. The remaining cells were loaded onto a Berkeley Lights Beacon optofluidic instrument in a novel plasma cell survival medium, and antigen reactivity of secreted antibody from individual B cells was measured for thousands of cells (**Figure S2**). Antigen-reactive B cells were exported from the instrument and the heavy and light chain genes from single B cells were sequenced and cloned into immunoglobulin expression vectors.

Using the parallel workflows, we isolated 386 recombinant SARS-CoV-2-reactive human mAbs that expressed sufficiently well as recombinant IgG to characterize the activity of the mAb. The recombinant mAbs were tested for binding in ELISA to recombinant monomeric S_RBD_ or S_NTD_ of SARS-CoV-2 or trimeric S2P_ecto_ proteins of SARS-CoV-2 or SARS-CoV (**Figure 3a**) and in a cell-impedance based SARS-CoV-2 neutralization assay with WA1/2020 strain SARS-CoV-2 in Vero-furin cells (**Figure S3**). The ELISA and neutralization screening assays revealed that the antibodies could be grouped into five binding patterns based on domain recognition and cross-reactivity (**Figure 3b)**. Comparison of binding patterns with full or partial neutralizing activity in a rapid cell-impedance-based SARS-CoV-2 neutralization test (**Figure 3c**) showed clearly that most of the neutralizing antibodies mapped to the RBD, revealing the RBD as the principal site of vulnerability for SARS-CoV-2 neutralization in these subjects. We examined the sequences for the 386 antibodies to assess the diversity of antigen-specific B cell clonotypes discovered. The analysis showed that among the 386 mAbs, 324 unique amino sequences were present and 311 unique V_H_-J_H_-CDRH3-V_L_-J_L_-CDRL3 clonotypes were represented, with diverse usage of antibody variable genes (**Figure 3d**). The CDR3 amino acid length distributions in the heavy and light chains were typical of human repertoires (**Figure 3e**)^23^. The high relatedness of sequences to the inferred unmutated ancestor antibody genes observed for this panel of antibodies (**Figure 3f**) contrasts with the much higher frequencies seen in B cell recall responses against common human pathogens like influenza^24^. These data suggest that the SARS-CoV-2 antibodies likely were induced during the primary response to SARS-CoV-2 infection and not by a recall response to a distantly related seasonal coronavirus. To validate the neutralizing activity measured in the cell-impedance-based neutralization test, we also confirmed neutralization for representative mAbs in a quantitative FRNT assay for SARS-CoV-2 (**Figure 3g**) or a neutralization assay with a SARS-CoV luciferase reporter virus (**Figure 3h**). Together, these results confirmed that mAbs recognizing multiple epitopes on S were able to neutralize SARS-CoV-2 and cross-react with SARS-CoV, with most neutralizing mAbs specific for the RBD of S.

**Figure 3.**
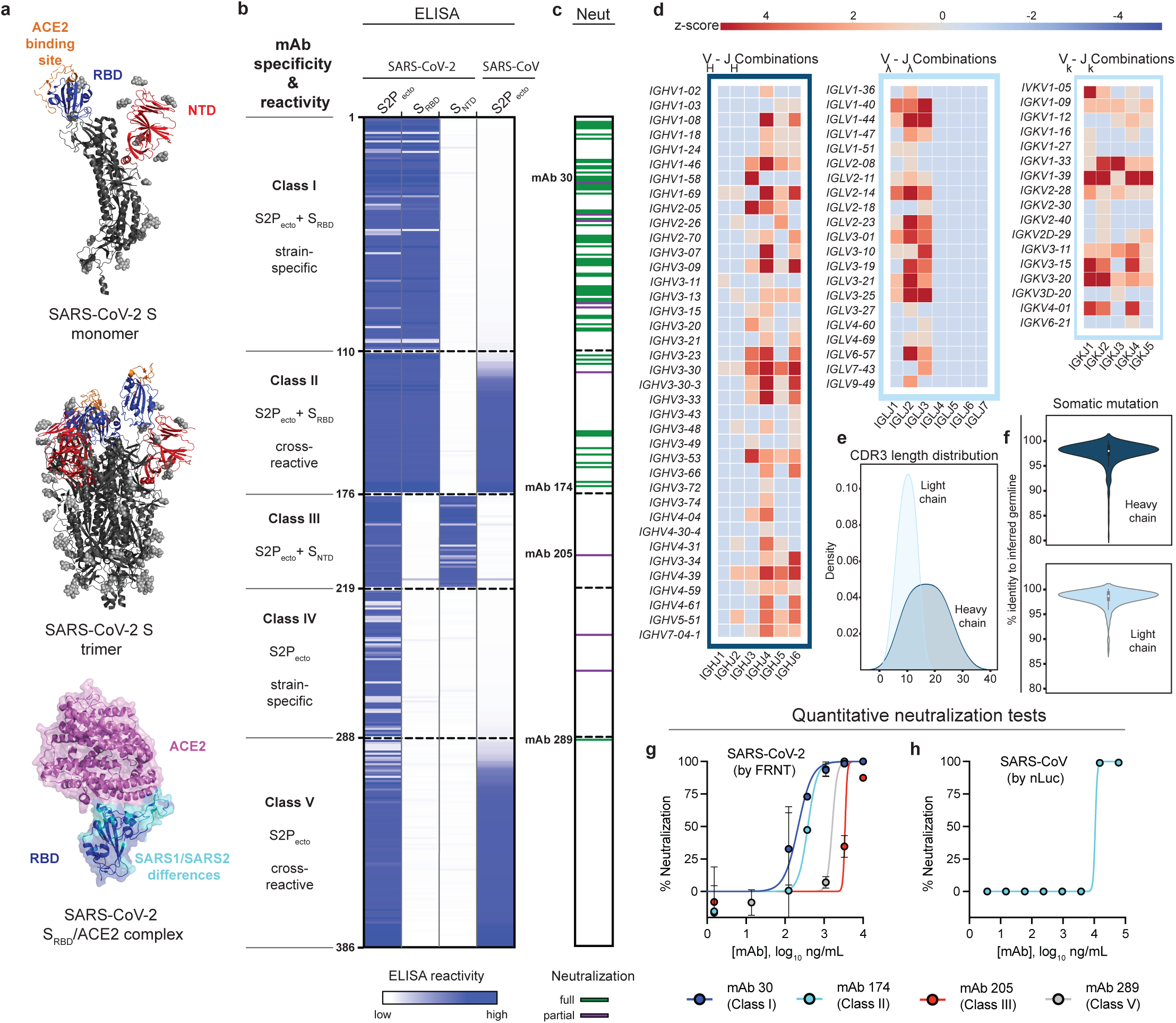
Reactivity and functional activity of 386 human mAbs. **a. Structures of SARS-CoV-2 spike antigen**. Top panel: S protein monomer of SARS-CoV-2 highlighting RBD (blue) and NTD (red) subdomains that were expressed as recombinant proteins. The ACE2 binding site on RBD is shown in orange. Known glycans are shown as light grey spheres. (PDB 6VYB) Middle panel: the structure of trimeric SARS-CoV-2 spike with one RBD in the “head up” conformation. Bottom panel: structure (PDB 6M0J) of SARS-CoV-2 RBD (blue) and ACE2 (pink) highlighting differences between RBDs of SARS-CoV-2 and SARS-CoV (cyan). **b. MAbs binding to each of four S proteins or subdomains**. The figure shows a heatmap for binding of 386 mAbs expressed recombinantly, representing optical density (O.D.) values collected at 450 nm for each antigen (range 0.035 to 4.5). White indicates lack of detectable binding, while blue indicates binding, with darker blue indicating higher O.D. values. **c. Screening test for neutralizing activity**. Each mAb was tested in a cell-impedance based neutralization test (**Figure S3**) using Vero-furin cells and live WA1/2020 strain SARS-CoV-2 in a BSL3 laboratory. Green indicates full protection of cells (full neutralization), purple indicates partial protection of cells, (partial neutralization), and white indicates neutralizing activity was not detected. Based on both binding and neutralization, we grouped the mAbs into classes. <u>Class I mAbs</u> bind to both S2P_ecto_ and S_RBD_ proteins and are SARS-CoV-2 specific; <u>Class II mAbs</u> also bind to both S2P_ecto_ and S_RBD_ proteins and cross-react with SARS-CoV; <u>Class III mAbs</u> bind to both S2P_ecto_ and S_NTD_ proteins and are mostly SARS-CoV-2 specific; <u>Class IV mAbs</u> bind only to S2P_ecto_ protein and are SARS-CoV-2 specific; <u>Class V mAbs</u> bind only to S2P_ecto_ protein and cross-react with SARS-CoV. **d. Heatmap showing usage of antibody variable gene segments for variable (V) and joining (J) genes**. Of the 386 antibodies tested in (**b**) and (**c**) above, 324 were found to have unique sequences, and those unique sequences were analyzed for genetic features. The frequency counts are derived from the total number of unique sequences with the corresponding V and J genes. The V/J frequency counts then were transformed into a z-score by first subtracting the average frequency, then normalizing by the standard deviation of each subject. Red denotes more common gene usage, while blue denotes less common gene usage. **e. CDR3 amino acid length distribution**. The CDR3 of each sequence was determined using PyIR software. The amino acid length of each CDR3 was counted. The distribution of CDR3 amino acid lengths for heavy or light chains then was plotted as a histogram and fitted using kernel density estimation for the curves. **f. Divergence from inferred germline gene sequences**. The number of mutations from each inferred unmutated ancestor sequence in the region spanning from antibody framework region 1 to 4 was counted up for each chain. These numbers then were transformed into percent values and plotted as violin plots. **g. Quantitative test for neutralizing activity against SARS-CoV-2 using FRNT**. Representative mAbs that exhibited full neutralizing activity in the screening neutralization test in (**c**) above using micro-scale expression were prepared in midi-scale as purified IgG and tested in a serial dilution series in the FRNT with live WA1/2020 strain SARS-CoV-2 to demonstrate neutralizing potency of class-representative mAbs. % neutralization of virus infection (relative to control wells with no mAb) at each dilution is shown. Values shown are the mean of two technical replicates, and error bars denote the standard deviation for each point. **h. Quantitative test for neutralizing activity against SARS-CoV using a nano-luciferase virus**. A representative purified mAb that exhibited cross-reactive binding to SARS-CoV S2P_ecto_ protein in (**b**) above and that also exhibited full neutralizing activity in the screening neutralization test in (**c**) above was tested in a serial dilution series in a neutralization test with a recombinant, reverse-genetics-derived SARS-CoV encoding a nano-luciferase reporter gene, and reduction of luciferase activity was used to calculate % neutralization. Values shown are the mean of two technical replicates, and error bars denote the standard deviation for each point.

Here, we coupled single-B-cell RNAseq methods with high-throughput IgG micro-expression and real-time neutralization assays to isolate and profile a large number of neutralizing antibodies in a period of only weeks after sample acquisition. As we show, recent advances in single-cell sequencing and gene synthesis have enabled antibody discovery at unprecedented scale and speed. In our particular example, sequences of confirmed neutralizing antibodies were transferred to downstream manufacturing partners only 18 days after antigen-specific cell sorting. However, given the need for affinity maturation and the development of a mature B cell response there are limits on the timeline from infection to isolation of potent neutralizing antibodies with therapeutic promise. It has been previously shown for Ebola virus infection that potently neutralizing antibodies are not easily isolated from memory B cells until later timepoints in the first year after infection^25,26^. It is likely that our success in isolating neutralizing antibodies from subjects 3 and 4 here at 50 days after onset, but not from subjects 1 and 2 at 35 or 36 days after onset, reflected additional maturation of the memory B cell response that occurred in the additional two weeks convalescence.

Overall, our work illustrates the promise of coupling recent technological advances for antibody discovery and defines the RBD of SARS-CoV-2 S as a major site of vulnerability for vaccine design and therapeutic antibody development. The most potent neutralizing human mAbs isolated here also could serve as candidate biologics to prevent or treat SARS-CoV-2 infection.

## Acknowledgements

We thank Merissa Mayo and Norma Suazo Galeano for coordination of human subjects studies, David O’Connor, Nasia Safdar, Geoff Baird, Jay Shendure and Samira Mubareka for helpful advice on human subjects, Angela Jones and the staff of the Vanderbilt VANTAGE core laboratory for expedited sequencing, Ross Trosseth for assistance with data management and analysis, Robin Bombardi and Cinque Soto for technical consultation on genomics approaches, Arthur Kim for production of a recombinant form of the mAb CR3022, Chris Swearingen and the staff of Fedex Express Specialty Services for expedited transport services, Vincent Pai and Keith Breinlinger of Berkeley Lights, Inc., and Kevin Louder and scientists at Twist Bioscience, Brian Fritz at 10x Genomics, and representatives at ACEA Biosciences for providing resources, outstanding expedited services and expert applications support. We thank Andrew Ward, Sandhya Bangaru and Nicole-Kallewaard-Lelay for protein reagents. This study was supported by Defense Advanced Research Projects Agency (DARPA) grant HR0011-18-2-0001, NIH contracts 75N93019C00074 and 75N93019C00062 and the Dolly Parton COVID-19 Research Fund at Vanderbilt. This work was supported by NIH grant 1S10RR028106-01A1 for the Next Generation Nucleic Acid Sequencer, housed in Vanderbilt Technologies for Advanced Genomics (VANTAGE) and supported by the National Center for Research Resources, Grant UL1 RR024975-01, and is now at the National Center for Advancing Translational Sciences, Grant 2 UL1 TR000445-06. S.J.Z. was supported by NIH T32 AI095202. J.B.C. is supported by a Helen Hay Whitney Foundation postdoctoral fellowship D.R.M. was supported by NIH T32 AI007151 and a Burroughs Wellcome Fund Postdoctoral Enrichment Program Award. J.E.C. is the recipient of the 2019 Future Insight Prize from Merck KGaA, Darmstadt Germany, which supported this research with a research grant. The content is solely the responsibility of the authors and does not necessarily represent the official views of the U.S. government or the other sponsors.

## Author contributions

Conceived of the project: S.J.Z., P.G., R.H.C., L.B.T., M.S.D., J.E.C.; Obtained funding: J.E.C. and M.S.D. Obtained human samples: M.O., H.Y.C., J.E.C.; Performed laboratory experiments: S.J.Z., P.G., R.E.C., J.X.R., A.T., R.S.N., R.E.S., N.S., E.B., J.E.D., K.W.M., S.S., D.R.M; Performed computational work: E.C.C., T.J., S.D., L.M.; Supervised research: M.S.D., L.B.T., R.S.B., R.H.C., J.E.C. Provided critical reagents: J.E.D., K.W.M., F.E.-H.L., D.C.N., I.S., R.S.B. Wrote the first draft of the paper: S.J.Z., P.G., R.H.C., J.E.C.; All authors reviewed and approved the final manuscript.

## Data availability

All relevant data are included with the manuscript.

## Supplementary Information

**Figure S1.**
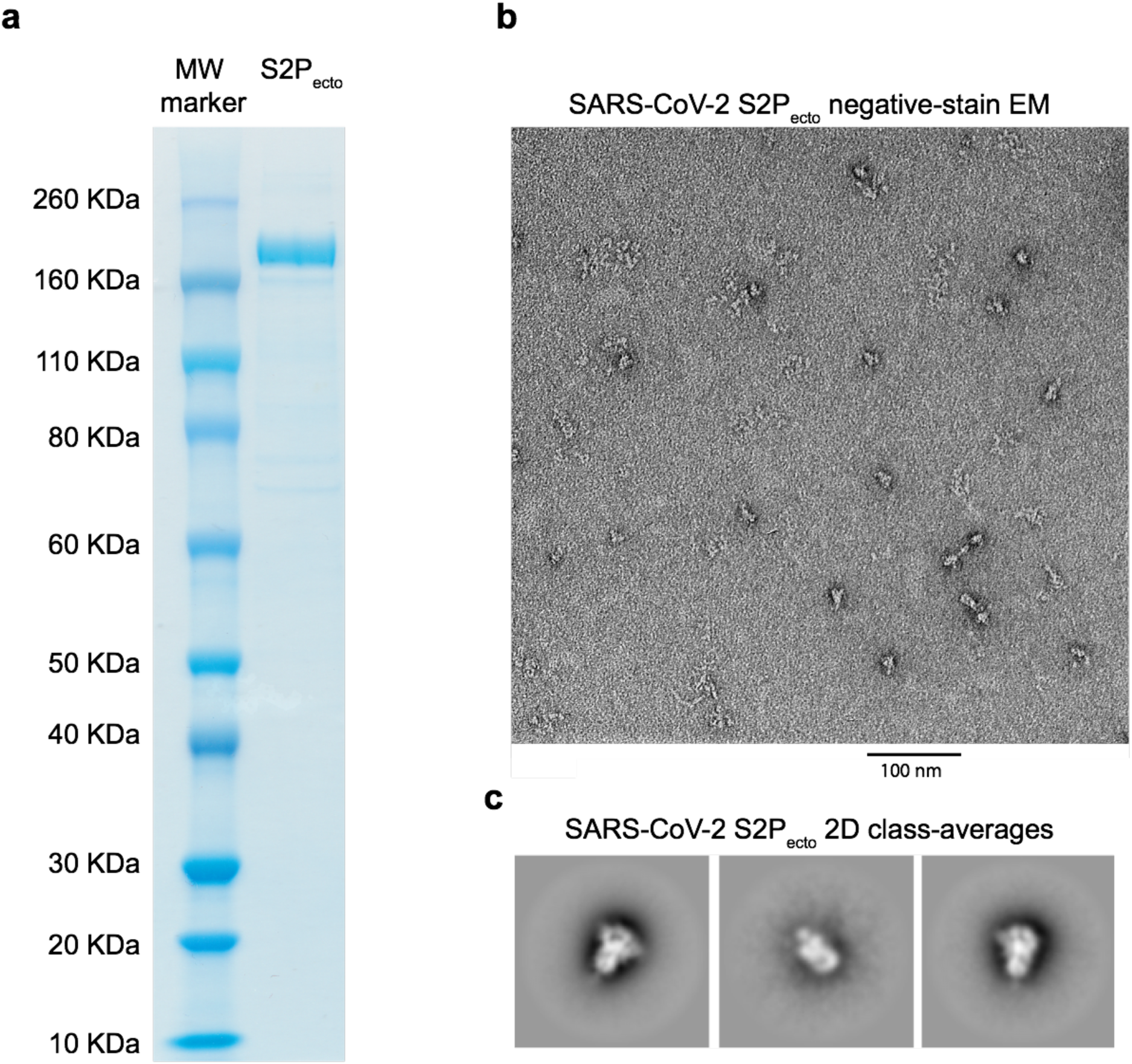
Expression and validation of prefusion-stabilized SARS-CoV-2 S2P_ecto]_]protein. **a**. Reducing SDS-PAGE gel indicating S2P_ecto_ protein migrating at approximately 180KDa. **b. Representative micrograph of negative-stain electron microscopy with S2P_ecto_ protein preparation**. Scale bar denotes 100 nm. **c. 2D class-averages of S2P_ecto_ protein in the prefusion conformation**. The size for each box is 128 pixels.

**Figure S2.**
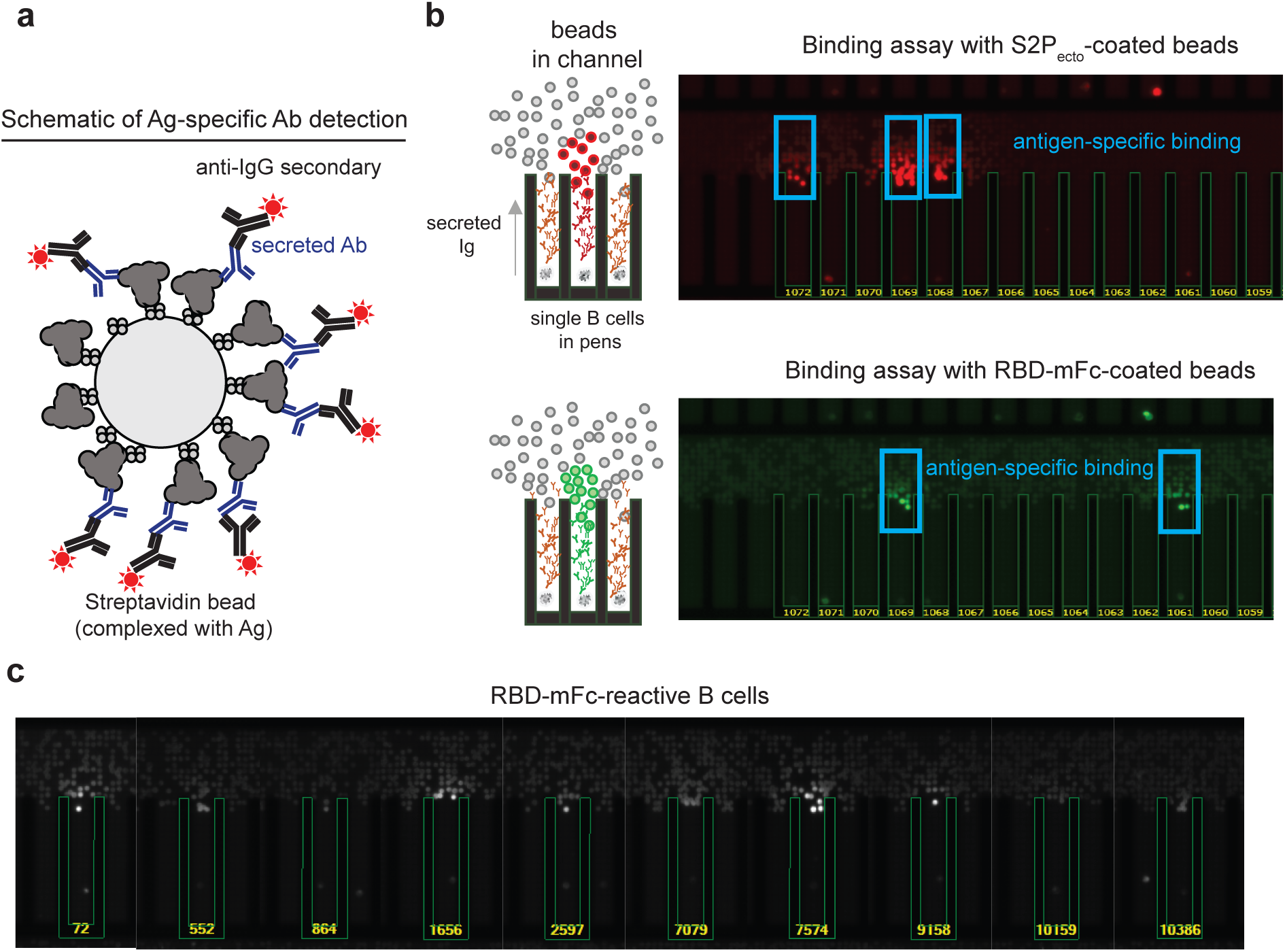
Functional assays from single antigen-reactive B cells. **a. Schematic of detection of antigen-specific antibody**. Biotinylated antigen (dark grey) was coupled to a streptavidin-conjugated polystyrene bead (light grey). Antibodies (blue) are secreted by single B cells loaded into individual NanoPens on the Berkeley Lights Beacon optofluidic device. Antibody binding to antigen was detected with a fluorescent anti-human IgG secondary Ab (black). **b. Left**: Schematic of fluorescing beads in the channel above a pen containing an individual B cell indicates antigen-specific reactivity. **Top right**: False-color still image of positive wells with B cells secreting S2P_ecto_-reactive antibodies. Reactive antibody diffusing out of a pen is visualized as a plume of fluorescence. **Bottom right**: False-color still image of positive wells with B cells secreting RBD-mFc-reactive antibodies. **c**. Representative images of RBD-mFc reactive clones.

**Figure S3.**
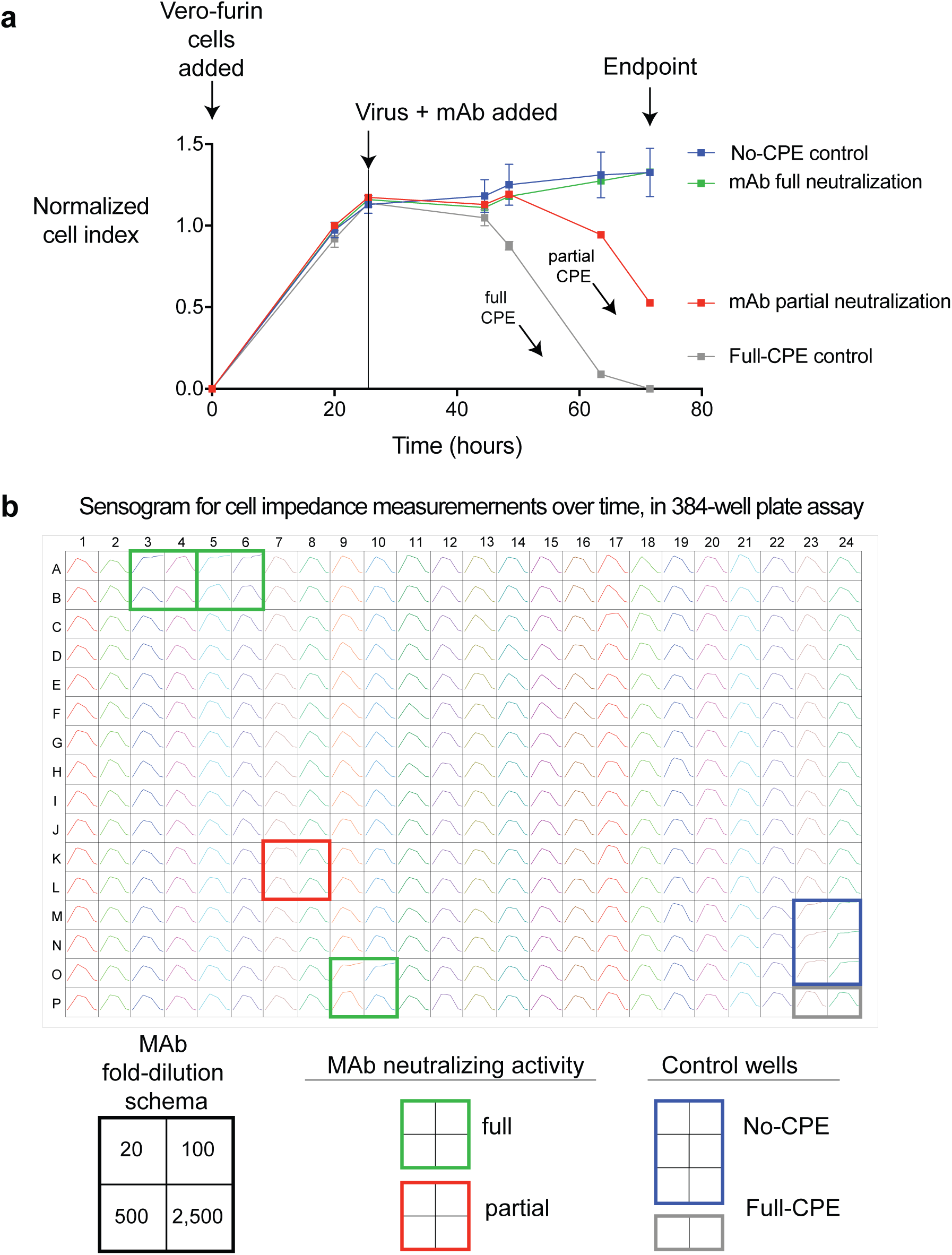
Real-time cell analysis assay to screen for neutralization activity. **a. Representative sensograms for neutralizing mAbs.** Curves for fully neutralizing mAb (green) and partially neutralizing mAb (red) by monitoring of CPE in Vero-furin cells that were inoculated with SARS-CoV-2 and pre-incubated with the respective mAb. Uninfected cells (blue) and infected cells without antibody addition (grey) served as controls for intact monolayer and full CPE, respectively. Data represented single well measurement for each mAb and mean SD values of technical duplicates or quadruplicates for the controls. **b. Example sensograms from individual wells of 384-well E-plate analysis showing rapid identification of SARS-CoV-2 neutralizing mAbs.** Neutralization was assessed using micro-scale purified mAbs and each mAb was tested in four 5-fold dilutions as indicated. Plates were measured every 8-12 hours for a total of 72 hrs as in (**a**).

**Movie S1. Time-lapse imaging of antigen-sorted single B cells secreting S2P_ecto_-reactive antibodies**. A field of view of the optofluidic chip is shown with single B cells at the bottom of NanoPens, as in Figure S1. Antigen-reactive antibody bound to S2P_ecto_ antigen conjugated to streptavidin polystyrene beads loaded into the channel is detected by an anti-IgG secondary antibody. Positive wells are identified by the specific bloom of fluorescence signal, indicating antigen-specific antibody diffusing out of a single pen. The edges of pens are highlighted in green and pen numbers are shown in yellow. For that field of view, there were 96 pens containing B cells, with 53 cells secreting trimer-reactive antibody and 30 cells secreting antibody reactive to RBD. The movie is composed of still images obtained every five minutes over the course of a 30-minute assay.

## Supplemental Tables

**Table S1.**
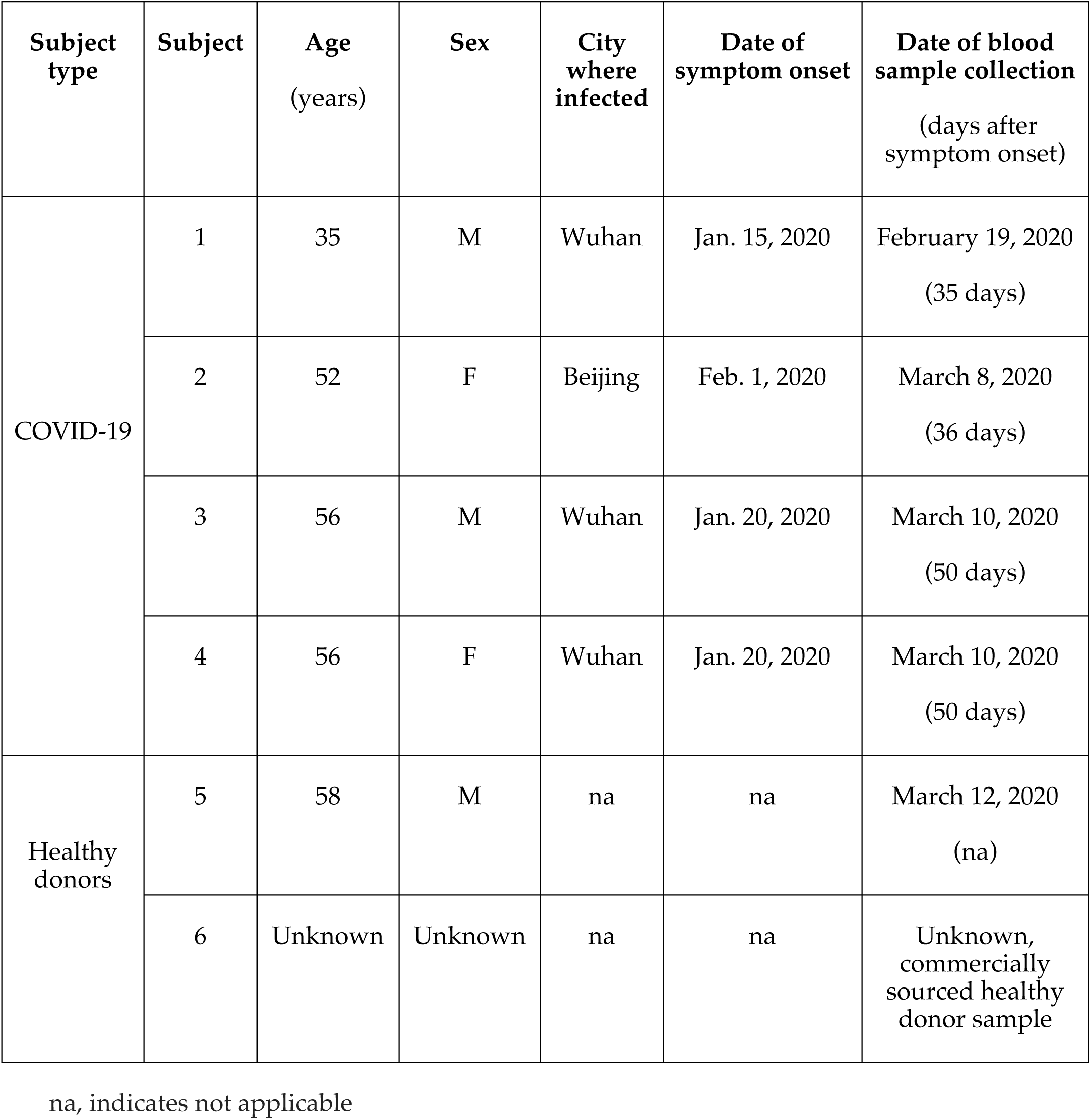
Human subjects studied in this paper

**Table S2.** Workflows used to isolate SARS-CoV-2 specific human mAbs from single B cells

| Single B cell technique<br>(number of Ag-reactive mAbs recovered) | Ag used for cell sorting | Feeder layer stimulation | Functional screen on instrument | Sequencing technique<br>(cDNA used) | Expression construct |
| --- | --- | --- | --- | --- | --- |
| Beacon<br>(n = 76) | S2P<sub>ecto</sub> +/- S<sub>RBD</sub> | + | S2P<sub>ecto</sub> binding | PacBio SMRT<br>(PCR amplicon) | Directly cloned cDNA in pMCis_G1 vector |
|  |  |  | RBD-mFc binding |  |  |
|  |  |  | RBD-mFc binding + ACE2 blocking |  |  |
| Chromium<br>(n = 310) | S2P<sub>ecto</sub> | + | na | Illumina Novaseq<br>(Chromium libraries) | Synthesized cDNA in pTwist-mCis_G1 vector |
|  |  | - |  |  |  |
|  | S<sub>RBD</sub> | + |  |  |  |
|  |  | - |  |  |  |

**Table S3.**
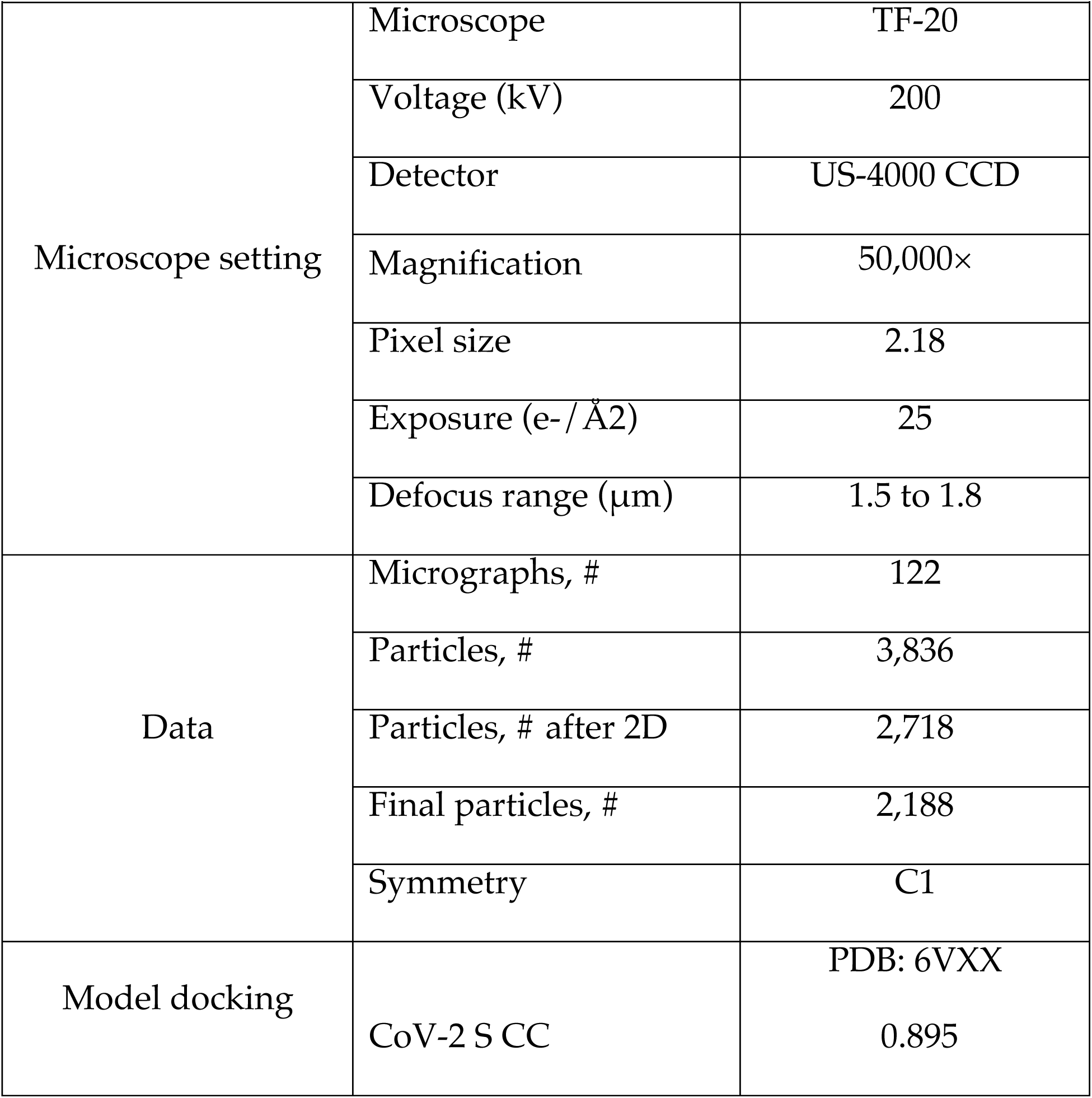
Summary of electron microscopy data collection and statistics SARS-CoV-2 S2P_ecto_ protein.

|  |  |  |
| --- | --- | --- |
| Microscope setting | Microscope | TF-20 |
|  | Voltage (kV) | 200 |
|  | Detector | US-4000 CCD |
|  | Magnification | 50,000× |
|  | Pixel size | 2.18 |
|  | Exposure (e-/Å<sup>2</sup>) | 25 |
|  | Defocus range (μm) | 1.5 to 1.8 |
| Data | Micrographs, # | 122 |
|  | Particles, # | 3,836 |
|  | Particles, # after 2D | 2,718 |
|  | Final particles, # | 2,188 |
|  | Symmetry | C1 |
| Model docking |  | PDB: 6VXX |
|  | CoV-2 S CC | 0.895 |

## Online Methods

### Research subjects

We studied four subjects in North America with recent laboratory-confirmed symptomatic SARS-CoV-2 infection that were acquired in China (**Table S1)**. The studies were approved by the Institutional Review Board of Vanderbilt University Medical Center, and subsite studies were approved by the Institutional Review Board of the University of Washington or the Research Ethics Board of the University of Toronto. Samples were obtained after written informed consent. Subject 1 (35-year-old male) was the earliest reported case of SARS-CoV-2 infection in the U.S. who presented with disease in Seattle, WA on January 19, 2020^1^, a blood sample was obtained for study on February 19, 2020. Subject 2 (52-year-old female) was infected following close exposure in Beijing, China to an infected person from Wuhan, China during the period between January 23 to January 29, 2020. She presented with mild respiratory disease symptoms from February 1 to 4, 2020 that occurred after travel to Madison, Wisconsin, USA. She obtained a diagnosis of infection by testing at the U.S. Centers for Disease Control on February 5, 2020. Blood samples were obtained for study on March 7 and March 8, 2020. Subject 3 (a 56-year-old male) and subject 4 (a 56-year-old female) are a married couple and residents of Wuhan, China who traveled to Toronto, Canada on January 22, 2020. Subject 3 first developed a cough without fever on January 20, 2020 in the city of Wuhan, where he had a normal chest x-ray on that day. He flew to Canada with persisting cough and arrived in Canada January 22, 2020 where he became febrile. He presented to a hospital in Toronto, January 23, 2020 complaining of fever, cough and shortness of breath; a nasopharyngeal swab was positive by PCR testing for SARS-CoV-2. His chest x-ray at that time was abnormal, and he was admitted for non-ICU impatient care. He improved gradually with supportive care, was discharged January 30, 2020 and rapidly became asymptomatic except for a residual dry cough that persisted for a month. He had a negative nasopharyngeal swab PCR test on February 19, 2020. Subject 4 is the wife of subject 3 who traveled with her husband from Wuhan. She was never symptomatic with respiratory symptoms or fever but was tested because of her exposure. Her nasopharyngeal swab was positive for SARS-CoV-2 by PCR, on January 24, 2020; repeat testing in followup on February 21, 2020 was negative. PBMCs were obtained by leukopheresis from subjects 3 and 4 on March 10, 2020, which was 50 days since the symptom onset of subject 3. Samples were transferred to Vanderbilt University Medical Center in Nashville, TN, USA on March 14, 2020.

### Cell culture

Vero E6 (CRL-1586, American Type Culture Collection (American Type Culture Collection, ATCC), Vero CCL81 (ATCC), HEK293 (ATCC), and HEK293T (ATCC) were maintained at 37°C in 5% CO_2_ in Dulbecco’s minimal essential medium (DMEM) containing 10% (vol/vol) heat-inactivated fetal bovine serum (FBS), 10 mM HEPES pH 7.3, 1 mM sodium pyruvate, 1× non-essential amino acids, and 100 U/mL of penicillin–streptomycin. Vero-furin cells were obtained from T. Pierson (NIH) and have been described previously^2^. Expi293F cells (ThermoFisher Scientific, A1452) were maintained at 37°C in 8% CO_2_ in Expi293F Expression Medium (ThermoFisher Scientific, A1435102). ExpiCHO cells (ThermoFisher Scientific, A29127) were maintained at 37°C in 8% CO_2_ in ExpiCHO Expression Medium (ThermoFisher Scientific, A2910002). Mycoplasma testing of Expi293F and ExpiCHO cultures was performed on a monthly basis using a PCR-based mycoplasma detection kit (ATCC, 30-1012K).

### Viruses

SARS-CoV-2 strain 2019 n-CoV/USA_WA1/2020 was obtained from the Centers for Disease Control and Prevention (a gift from Natalie Thornburg). Virus was passaged in Vero CCL81 cells and titrated by plaque assay on Vero E6 cells. All work with infectious SARS-CoV-2 was approved by the Washington University School of Medicine or UNC-Chapel Hill Institutional Biosafety Committees and conducted in approved BSL3 facilities using appropriate powered air purifying respirators and personal protective equipment.

### Recombinant antigens and proteins

A gene encoding the ectodomain of a prefusion conformation-stabilized SARS-CoV-2 spike (S2P_ecto_) protein was synthesized and cloned into a DNA plasmid expression vector for mammalian cells. A similarly designed S protein antigen with two prolines and removal of the furin cleavage site for stabilization of the prefusion form of S was reported previously^3^. Briefly, this gene includes the ectodomain of SARS-CoV-2 (to residue 1,208), a T4 fibritin trimerization domain, an AviTag site-specific biotinylation sequence, and a C-terminal 8x-His tag. To stabilize the construct in the prefusion conformation, we included substitutions K968P and V969P and mutated the furin cleavage site at residues 682-685 from RRAR to ASVG. This recombinant spike 2P-stabilized protein (designated here as S2P_ecto_) was isolated by metal affinity chromatography on HisTrap Excel columns (GE Healthcare), and protein preparations were purified further by size-exclusion chromatography on a Superose 6 Increase 10/300 column (GE Healthcare). The presence of trimeric, prefusion conformation S protein was verified by negative-stain electron microscopy (**Figure S1)**. To express the S_RBD_ subdomain of SARS-CoV-2 S protein, residues 319-541 were cloned into a mammalian expression vector downstream of an IL-2 signal peptide and upstream of a thrombin cleavage site, an AviTag, and a 6x-His tag. RBD protein fused to mouse IgG1 Fc domain (designated RBD-mFc), was purchased from Sino Biological (40592-V05H). For B cell labeling and sorting, RBD-mFc and S2P_ecto_ proteins were biotinylated using the EZ-Link™ Sulfo-NHS-LC-Biotinylation Kit and vendor’s protocol (ThermoFisher Scientific, 21435).

### Electron microscopy (EM) stain grid preparation and imaging of SARS-CoV-2 S2P_ecto_ protein

For screening and imaging of negatively-stained (NS) SARS-CoV-2 S2P_ecto_ protein, approximately 3 µL of the sample at concentrations of about 10 to 15 µg/mL was applied to a glow discharged grid with continuous carbon film on 400 square mesh copper EM grids (Electron Microscopy Sciences, Hatfield, PA). The grids were stained with 0.75% uranyl formate (UF)^4^. Images were recorded on a Gatan US4000 4k × 4k CCD camera using an FEI TF20 (TFS) transmission electron microscope operated at 200 keV and control with SerialEM^5^. All images were taken at 50,000× magnification with a pixel size of 2.18 Å/pix in low-dose mode at a defocus of 1.5 to 1.8 μm. Total dose for the micrographs was ∼25 e^−^/Å^2^. Image processing was performed using the cryoSPARC software package^6^. Images were imported, and particles were CTF estimated. The images then were denoised and picked with Topaz^7^. The particles were extracted with a box size of 256 pixels and binned to 128 pixels. 2D class averages were performed and good classes selected for *ab-initio* model and refinement without symmetry (see also **Table S3** for details). For EM model docking of SARS-CoV-2 S2P_ecto_ protein, the closed model (PDB: 6VXX) was used in Chimera^8^ for docking to the EM map (see also **Table S3** for details).

### Human subject selection and target-specific memory B cells isolation

B cell responses to SARS-CoV-2 in PBMCs from a cohort of four subjects with documented previous infection with the virus were analyzed for antigen specificity, and PBMCs were used for SARS-CoV-2-specific B cell enrichment. The frequency of SARS-CoV-2 S protein-specific B cells was identified by antigen-specific staining with either biotinylated S2P_ecto_ or RBD-mFc protein.

Briefly, B cells were purified magnetically (STEMCELL Technologies) and stained with anti-CD19-APC (clone HIB19, 982406), -IgD-FITC (clone LA6-2, 348206), and -IgM-FITC (clone MNM-88, 314506) phenotyping antibodies (BioLegend) and biotinylated antigen. A 4′,6-diamidino-2-phenylindole (DAPI) stain was used as a viability dye to discriminate dead cells. Antigen-labeled class-switched memory B cell-antigen complexes (CD19^+^IgM^-^IgD^-^Ag^+^DAPI^-^) were detected with a R-phycoerythrin (PE)-labeled streptavidin conjugate (ThermoFisher Scientific, S866). After identification of the two subjects with the highest B cell response against SARS-CoV-2 (subjects 3 and 4), target-specific memory B cells were isolated by flow cytometric sorting using an SH800 cell sorter (Sony) from pooled PBMCs of these two subjects, after labeling of B cells with either biotinylated S2P_ecto_ or RBD-mFc proteins.

Overall, from > 4 x 10^8^ PBMCs, 2,126 RBD-mFc-reactive and 5,544 S2P_ecto_ protein-reactive B cells were sorted and subjected to further analysis. Several methods were implemented for the preparation of sorted B cells for sequencing. Approximately 4,500 sorted cells were subjected to direct sequencing immediately after flow cytometric sorting. The remaining cells were expanded in culture for eight days in the presence of irradiated 3T3 feeder cells that were engineered to express human CD40L, IL-21, and BAFF, as described previously^9^. The expanded lymphoblastoid cell lines (LCLs) secreted high levels of S protein-specific antibodies, as confirmed by ELISA to detect antigen-specific human antibodies in culture supernatants. Approximately 40,000 expanded LCLs were sequenced using the Chromium sequencing method (10x Genomics).

### Microfluidic device selection of single antigen-specific B cells

Activated memory B cells were screened using Berkeley Lights’ Beacon®™ optofluidic system. Purified B cell samples were imported automatically onto OptoSelect™ 11k chips in a novel plasmablast survival medium that promotes antibody secretion and preserves cell viability^10^. Single-cell penning was then performed using OEP™ technology in which light is used to transfer B cells into individual nanoliter-volume chambers (NanoPens™). Using this light-based manipulation, thousands of LCLs were transferred into pens across multiple chips in each workflow. We performed an on-chip, fluorescence-based assay to identify antibodies that bound SARS-CoV-2 S2P_ecto_ or RBD-mFc protein. We prepared 6-to 8-micron and 10-to 14-micron RBD-mFc-conjugated beads by coupling biotinylated RBD-mFc protein to streptavidin-coated polystyrene particles (Spherotech Inc.). We prepared 6-to 8-micron S2P_ecto_ protein-conjugated beads by coupling full-length S2P_ecto_ protein to streptavidin-coated polystyrene particles.

Assays consisted of mixing beads conjugated with the RBD-mFc or S2P_ecto_ proteins with fluorescently-labeled anti-human secondary antibodies (AF568, Thermo Fisher Scientific) and importing this assay mixture into OptoSelect 11k chips. Antigen-specific antibodies bound the antigen-conjugated beads, which then sequestered the fluorescent secondary antibody. Cells secreting antigen-specific antibodies were identified by locating the NanoPens immediately adjacent to the fluorescent beads. Antigen-specific cells of interest were exported from specific NanoPen chambers to individual wells of 96-well RT-PCR plates containing lysis buffer.

### Sequencing and cloning of single antigen-specific B cells

After export from the Beacon, antibody heavy and light chain sequences for B cells secreting antibodies with RBD-mFc-or S2P_ecto_-binding antibodies were amplified and recovered using components of the Opto™ Plasma B Discovery cDNA Synthesis Kit (Berkeley Lights). Antibody heavy and light chain sequences were amplified through a 5’RACE approach using the kit’s included “BCR Primer 2” forward primer and isotype-specific reverse primers. The 5’-RACE amplified cDNA was sequenced using the Pacific Biosciences Sequel platform using the SMRTbell Barcoded Adapt Complete Prep-96 kit (Pacific Biosciences) and a 6-hour movie time. In a redundant sequencing approach, heavy and light chain sequences were amplified using a cocktail of custom V and J gene-specific primers (similar to previously described human Ig gene-specific primers^11^) from the original 5’-RACE-amplified cDNA while the products of the gene-specific amplification were sent for Sanger sequencing (GENEWIZ). The sequences generated by these two approaches were analyzed using our Python-based antibody variable gene analysis tool (PyIR; https://github.com/crowelab/PyIR)^12^ to identify which V and J genes most closely matched the nucleotide sequence. Heavy and light chain sequences were then amplified from the original cDNA using cherry-picked V and J gene-specific primers most closely corresponding to the V and J gene calls made by PyIR. These primers include adapter sequences which allow Gibson-based cloning into a monocistronic IgG1 expression vector (pMCis_G1). Similar to a vector described below, this vector contains an enhanced 2A sequence and GSG linker that allows simultaneous expression of mAb heavy and light chain genes from a single construct upon transfection^13^. The pMCis_G1 vector was digested using the New England BioLabs restriction enzyme FspI, and the amplified paired heavy and light chain sequences were cloned through Gibson assembly using NEBuilder HiFi DNA Assembly Master Mix. After recovered sequences were cloned into pMCis_G1 expression constructs, recombinant antibodies were expressed in Chinese hamster ovary (CHO) cells and purified by affinity chromatography as detailed below. Antigen-binding activity was confirmed using plate-based ELISA.

### Generation of single-cell antibody variable genes profiling libraries

As an alternative approach, we also used a second major approach for isolation of SARS-CoV-2-reactive antibodies. In some experiments, the Chromium Single Cell V(D)J workflow with B-cell only enrichment option was used for generating linked heavy-chain and light-chain antibody profiling libraries. Approximately 2,866 directly sorted S2P_ecto_ or 1,626 RBD-mFc protein-specific B cells were split evenly into two replicates each and separately added to 50 μL of RT Reagent Mix, 5.9 μL of Poly-dt RT Primer, 2.4 μL of Additive A and 10 μL of RT Enzyme Mix B to complete the Reaction Mix as per the vendor’s protocol, which then was loaded directly onto a Chromium chip (10x Genomics). Similarly, for the remaining sorted cells that were expanded in bulk, approximately 40,000 cells from two separate sorting approaches were split evenly across four reactions and processed separately as described above before loading onto a Chromium chip. The libraries were prepared following the User Guide for Chromium Single Cell V(D)J Reagents kits (CG000086_REV C).

### Next generation DNA sequence analysis of antibody variable genes

Chromium Single Cell V(D)J B-Cell enriched libraries were quantified, normalized and sequenced according to the User Guide for Chromium Single Cell V(D)J Reagents kits (CG000086_REV C). The two enriched libraries from direct flow cytometric cell sorting were sequenced on a NovaSeq sequencer (Illumina) with a NovaSeq 6000 S1 Reagent Kit (300 cycles) (Illumina). The four enriched libraries from bulk expansion were sequenced on a NovaSeq sequencer with a NovaSeq 6000 S4 Reagent Kit (300 cycles (Illumina). All enriched V(D)J libraries were targeted for sequencing depth of at least 5,000 raw read pairs per cell. Following sequencing, all samples were demultiplexed and processed through the 10x Genomics Cell Ranger software (version 2.1.1) as below.

### Bioinformatics analysis of single-cell sequencing data

The down-selection to identify lead candidates for expression was carried out in two phases. In the first phase, all paired antibody heavy and light chain variable gene cDNA nucleotide sequences obtained that contained a single heavy and light chain sequence were processed using PyIR. We considered heavy and light chain encoding gene pairs productive and retained them for additional downstream processing if they met the following criteria: 1) did not contain a stop codon, 2) encoded an intact CDR3 and 3) contained an in-frame junctional region. The second phase of processing eliminated redundant sequences (those with identical amino acid sequences). Any antibody variant that was designated as an IgM isotype (based on the sequence and assignment using the 10x Genomics Cell Ranger V(D)J software [version 2.1.1]) was removed from consideration (while IgG and IgA isotype antibodies were retained). The identities of antibody variable gene segments, CDRs, and mutations from inferred germline gene segments were determined by using PyIR.

### Antibody gene synthesis

Sequences of selected mAbs were synthesized using a rapid high-throughput cDNA synthesis platform (Twist Bioscience) and subsequently cloned into an IgG1 monocistronic expression vector (designated as pTwist-mCis_G1) for mammalian cell culture mAb secretion. This vector contains an enhanced 2A sequence and GSG linker that allows simultaneous expression of mAb heavy and light chain genes from a single construct upon transfection^13^.

### MAb production and purification

For high-throughput production of recombinant mAbs, we adopted approaches designated as “micro-scale” or “midi-scale”. For “micro-scale” mAbs expression, we performed micro-scale transfection (∼1 mL per antibody) of CHO cell cultures using the Gibco™ ExpiCHO™ Expression System and a protocol for deep 96-well blocks (ThermoFisher Scientific) as detailed in accompanying manuscript^14^. Briefly, synthesized antibody-encoding lyophilized DNA was reconstituted in OptiPro serum-free medium (OptiPro SFM) and used for transfection of ExpiCHO cell cultures into 96-deep-well blocks. For high-throughput micro-scale mAbs purification, clarified culture supernatants were incubated with MabSelect SuRe resin (Cytiva, formerly GE Healthcare Life Sciences), washed with PBS, eluted, buffer-exchanged into PBS using Zeba™ Spin Desalting Plates (Thermo Fisher Scientific) and stored at 4°C until use. For “midi-scale” mAbs expression, we performed transfection (∼15 mL per antibody) of CHO cell cultures using the Gibco™ ExpiCHO™ Expression System and protocol for 50 mL mini bioreactor tubes (Corning) as described by the vendor. For high-throughput midi-scale mAb purification, culture supernatants were purified using HiTrap MabSelect SuRe (Cytiva, formerly GE Healthcare Life Sciences) on a 24-column parallel protein chromatography system (Protein BioSolutions). Purified mAbs were buffer-exchanged into PBS, concentrated using Amicon® Ultra-4 50KDa Centrifugal Filter Units (Millipore Sigma) and stored at 4°C until use.

### ELISA binding screening assays

Wells of 96-well microtiter plates were coated with purified recombinant SARS-CoV-2 S protein, SARS-CoV-2 S_RBD_ protein, SARS-CoV-2 S_NTD_ (kindly provided by Nicole Kallewaard-Lelay, Astra Zeneca) or SARS-CoV S protein (kindly provided by Sandhya Bangaru and Andrew Ward, The Scripps Research Institute) at 4°C overnight. Plates were blocked with 2% non-fat dry milk and 2% normal goat serum in DPBS containing 0.05% Tween-20 (DPBS-T) for 1 hr. For mAb screening assays, CHO cell culture supernatants or purified mAbs were diluted 1:20 in blocking buffer, added to the wells, and incubated for 1 hr at ambient temperature. The bound antibodies were detected using goat anti-human IgG conjugated with HRP (horseradish peroxidase) (Southern Biotech) and TMB (3,3′,5,5′-tetramethylbenzidine) substrate (Thermo Fisher Scientific). Color development was monitored, 1N hydrochloric acid was added to stop the reaction, and the absorbance was measured at 450 nm using a spectrophotometer (Biotek). For dose-response assays, serial dilutions of purified mAbs were applied to the wells in triplicate, and mAb binding was detected as detailed above. Half-maximal effective concentration (EC_50_) values for binding were determined using Prism v8.0 software (GraphPad) after log transformation of mAb concentration using sigmoidal dose-response nonlinear regression analysis.

### Real-time cell analysis assay (RTCA)

To screen for neutralizing activity in the panel of recombinantly expressed mAbs, we used a high-throughput and quantitative real-time cell analysis (RTCA) assay and xCelligence RTCA HT Analyzer (ACEA Biosciences Inc.) that assesses kinetic changes in cell physiology, including virus-induced cytopathic effect (CPE). Twenty (20) μL of cell culture medium (DMEM supplemented with 2% FBS) was added to each well of a 96-well E-plate using a ViaFlo384 liquid handler (Integra Biosciences) to obtain background reading. Six thousand (6,000) Vero-furin cells in 20 μL of cell culture medium were seeded per each well, and the plate was placed on the analyzer. Sensograms were visualized using RTCA HT software version 1.0.1 (ACEA Biosciences Inc). For a screening neutralization assay, equal amounts of virus were mixed with micro-scale purified Abs in a total volume of 40 μL using DMEM supplemented with 2% FBS as a diluent and incubated for 1 hr at 37°C in 5% CO_2_. At ∼17-20 hrs after seeding the cells, the virus-mAb mixtures were added to the cells in 384-well E-plates. Wells containing virus only (in the absence of mAb) and wells containing only Vero cells in medium were included as controls. Plates were measured every 8-12 hours for 48 to 72 hrs to assess virus neutralization. Micro-scale antibodies were assessed in four 5-fold dilutions (starting from a 1:20 sample dilution), and their concentrations were not normalized. In some experiments mAbs were tested in triplicate using a single (1:20) dilution. Neutralization was calculated as the percent of maximal cell index in control wells without virus minus cell index in control (virus-only) wells that exhibited maximal CPE at 40 to 48 hrs after applying virus-antibody mixture to the cells. A mAb was classified as fully neutralizing if it completely inhibited SARS-CoV-2-induced CPE at the highest tested concentration, while a mAb was classified as partially neutralizing if it delayed but did not fully prevent CPE at the highest tested concentration. Representative sensograms for fully neutralizing and partially neutralizing mAbs are shown in Figure S3. For mAb potency ranking experiments, individual mAbs identified as fully neutralizing from the screening study were assessed by FRNT.

### Sequence analysis of antigen-reactive mAb sequences

Sequences of the 386 mAbs isolated by different approaches were combined and run through PyIR to identity the V genes, J genes, CDR3 lengths, and percent identity to germline, and sequence within the FR1-FR4 region for both heavy and light chains. Sequences were then deduplicated on the nucleotide sequences identified in the FR1-FR4 region. Among the 386 mAbs, there were 324 unique nucleotide sequences that were analyzed for V/D/J gene usage, CDR3 length, and somatic mutation. First, the number of sequences with corresponding V and J genes were counted. The V/J frequency counts were then transformed into a z-score by first subtracting away the average frequency then normalizing by the standard deviation of each subject. The z-score was then plotted as a heatmap using python seaborn library. The amino acid length of each CDR3 was counted. The distribution of CDR3 amino acid lengths were then plotted as histograms and fitted using kernel density estimation for the curves using python seaborn library. The number of mutations from each inferred germ-line sequence starting from FR-1 to FR4 was counted up for each chain. This number was then transformed into a percentage value. These values are then plotted as a categorical distribution plot as a violin plot using the python seaborn.catplot library.

### Focus reduction neutralization test (FRNT)

Serial dilutions of mAbs were incubated with 10^2^ FFU of SARS-CoV-2 for 1 hr at 37°C. The mAb–virus complexes were added to Vero E6 cell monolayers in 96-well plates for 1 hr at 37°C. Subsequently, cells were overlaid with 1% (w/v) methylcellulose in Minimum Essential Medium (MEM) supplemented to contain 2% heat-inactivated FBS. Plates were fixed 30 hrs later by removing overlays and fixed with 4% PFA in PBS for 20 min at room temperature. The plates were incubated sequentially with 1 µg/mL of rCR3022 anti-S antibody^15^ and horseradish-peroxidase (HRP)-conjugated goat anti-human IgG in PBS supplemented with 0.1% (w/v) saponin (Sigma) and 0.1% bovine serum albumin (BSA). SARS-CoV-2-infected cell foci were visualized using TrueBlue peroxidase substrate (KPL) and quantitated on an ImmunoSpot 5.0.37 Macro Analyzer (Cellular Technologies).

### SARS-CoV neutralization assays using SARS-CoV luciferase reporter virus

Full-length viruses expressing luciferase were designed and recovered via reverse genetics and described previously.^16,17^ Viruses were titered in Vero E6 USAMRID cell culture monolayers to obtain a relative light units (RLU) signal of at least 20X the cell-only control background. Vero E6 USAMRID cells were plated at 20,000 cells per well the day prior in clear-bottom black-walled 96-well plates (Corning #3904). Neutralizing antibodies were diluted serially 4-fold up to eight dilution times. SARS-Urbani NanoLuc virus was mixed with serially diluted antibodies. Antibody-virus complexes were incubated at 37°C in 5% CO_2_ for 1 hr. Following incubation, growth medium was removed and virus-antibody dilution complexes were added to the cells in duplicate. Virus-only and cell-only controls were included in each neutralization assay plate.

Following infection, plates were incubated at 37°C in 5% CO_2_ for 48 hours. After the 48-hour incubation, cells were lysed and luciferase activity was measured using the Nano-Glo Luciferase Assay System (Promega), according to the manufacturer’s specifications.

### High-throughput mAb quantification

High-throughput quantification of micro-scale produced mAbs was performed from CHO culture supernatants or micro-scale purified mAbs in a 96-well plate format using the Cy-Clone Plus Kit and an iQue Plus Screener flow cytometer (IntelliCyt Corp), according to the vendor’s protocol. Purified mAbs were assessed at a single dilution (1:10 final, using 2 μL of purified mAb per reaction), and a control human IgG solution with known concentration was used to generate a calibration curve. Data were analyzed using ForeCyt software version 6.2 (IntelliCyt Corp).

### Quantification and statistical analysis

The descriptive statistics mean ± SEM or mean ± SD were determined for continuous variables as noted. Technical and biological replicates are described in the figure legends. Statistical analyses were performed using Prism v8.0 (GraphPad).

